# Prostaglandin receptor EP2 is a novel molecular target for high-risk neuroblastoma

**DOI:** 10.1101/2020.02.24.963108

**Authors:** Ruida Hou, Ying Yu, Davis T. Nguyen, Madison N. Sluter, Lexiao Li, Jun Yang, Jianxiong Jiang

## Abstract

As the third-most common type of cancers in infants and young children, neuroblastoma accounts for nearly 10% of all childhood cancers. Despite remarkable advances in tumor diagnosis and management during the past decades, the five-year survival rates for patients with high-risk neuroblastoma remain below 50%. Developing new therapies for this devastating type of childhood cancer is an urgent unmet need. Cyclooxygenase (COX) via synthesizing prostaglandin E2 (PGE_2_) promotes tumor cell proliferation, survival, migration and invasion, and fosters an inflammation-enriched microenvironment that can facilitate angiogenesis, immune evasion and treatment resistance. However, which downstream PGE_2_ receptor subtype – namely EP1, EP2, EP3 and EP4 – is directly involved in COX activity-promoted neuroblastoma growth remains elusive. Analyzing five major neuroblastoma patient datasets: Versteeg-88, Kocak-649, SEQC-498, Primary NRC-283, and Oberthuer-251, we show that COX-1/PGE_2_/EP2 signaling axis is highly associated with the aggressiveness of human neuroblastoma. A time-resolved fluorescence resonance energy transfer (TR-FRET) method reveals EP2 as the key Gα_s_-coupled receptor that mediates PGE_2_-initiated cAMP signaling in neuroblastoma cells with various risk factors. Taking advantage of novel, selective and bioavailable small-molecule antagonists that we recently developed to target the PGE_2_/EP2 signaling *in vivo*, we have demonstrated that pharmacological inhibition of the peripheral EP2 receptor substantially impairs the growth of human neuroblastoma xenografts and the associated angiogenesis in mice. Collectively, our results suggest that the PGE_2_/EP2 pathway contributes to the growth and malignant potential of human neuroblastoma cells; pharmacological inhibition on EP2 receptor by our drug-like compounds might provide a novel therapeutic strategy for this deadly pediatric cancer.

## INTRODUCTION

Neuroblastoma is the most common extracranial solid tumor in childhood and accounts for 7-10% of all pediatric cancers, causing approximately 15% of all cancer-related deaths in young children. Owing to the tremendous progress in tumor diagnosis and management, the survival of children with neuroblastoma has much improved during the past decade. However, patients with high-risk neuroblastoma still have very poor prognosis with a five-year survival rate less than 50% even after surgery along with intensive chemotherapy, radiotherapy, stem cell transplantation, and immunotherapy (1). High-risk and poor-prognosis neuroblastoma in younger children is often associated with the amplification of the MYC family member, *MYCN*; whereas the deletion of the long arm of chromosome 11 (11q) that causes the development of aggressive neuroblastoma in older children often correlates with advanced disease stage, drug resistance, and decreased survival probability (2,3). Therefore, identifying new druggable targets and developing novel therapies for patients suffering from neuroblastoma – particularly those who are in high-risk category and developed resistance to current treatment – are in urgent demand.

As an essential component of innate immune system, inflammation in the acute phase represents a protective strategy of the body to remove the injurious stimuli and to initiate the healing process. However, inflammation often persists and becomes chronic, and prolonged severe inflammation is frequently associated with the occurrence and progression of a variety types of tumors including those of blood, brain, breast, colon, lung, pancreas, prostate, and skin (4–6). As such, cyclooxygenase (COX), a conventional inflammatory executor, has recently been demonstrated to provide important driving force for neuroblastoma pathogenesis (7). Daily oral intake of COX-selective inhibitors such as aspirin or diclofenac can decrease the burden of tumors with *MYCN*-amplification, reduce the presence of tumor-associated innate immune cells (8), and delay the rapid progression of 11q-deletion high-risk neuroblastoma (9). COXs – with two isoforms COX-1 and COX-2 – are the rate-limiting enzymes in the synthesis of bioactive lipids such as prostaglandin E2 (PGE_2_), which is often highly present in tumor tissues and can increase tumor aggressiveness (10). To synthesize PGE_2_, arachidonic acid first is converted by COX-1 or COX-2 to prostaglandin H2 (PGH_2_). Short-lived PGH_2_ then is further quickly catalyzed to PGE_2_ by tissue-specific isomerases – prostaglandin E synthases (PGESs) comprising microsomal prostaglandin E synthase-1 (mPGES-1), mPGES-2 and cytosolic PGES (cPGES) (11). Among the three PGES isozymes, mPGES-1 is considered inducible and functionally coupled to COX-2 (12). A recent study shows that the pharmacological inhibition of mPGES-1 suppressed 11q-deletion neuroblastoma tumor growth (13), attesting to an essential role for PGE_2_-mediated pro-inflammatory signaling in the development and progression of high-risk neuroblastoma.

PGE_2_ was demonstrated to regulate tumorigenesis through four G protein-coupled receptors (GPCRs), EP1, EP2, EP3 and EP4 (10,14–16), all of which were detected in human neuroblastoma cells and tissues (13,17). EP1 receptor is Gα_q_-coupled to mediate the mobilization of cytosolic Ca^2+^ and activation of protein kinase C (PKC); both EP2 and EP4 are linked to Gα_s_ activating adenylyl cyclase to generate cAMP; EP3 receptor is mainly coupled to Gα_i_, and its activation downregulates the cAMP signaling (15). To date, the EP subtype underlying the COX/PGES/PGE_2_ pathway-driven tumor growth in neuroblastoma remains elusive. Taking advantage of several novel bioavailable small-molecule compounds that we recently developed to suppress the chronic phase of inflammation in the periphery, the main objective of the present work was to identify the PGE_2_ receptor subtype that is involved in the pathogenesis of neuroblastoma as a novel molecular target. We also determined the therapeutic effects of these EP2-targeting and drug-like antagonists in a mouse model of high-risk neuroblastoma.

## MATERIALS AND METHODS

### Cell culture

Human neuroblastoma cell lines SK-N-SH (ATCC #: HTB-11), SK-N-BE(2) (ATCC #: CRL-2271), BE(2)-C (ATCC #: CRL-2268), SH-SY5Y(ATCC #: CRL-2266), SK-N-AS (ATCC #: CRL-2137), and human normal fibroblast cell line HS68 (ATCC #: CRL-1635) were cultured in Dulbecco’s modified Eagle’s medium (DMEM, Gibco) supplemented with 10% (v/v) fetal bovine serum (FBS, HyClone) and penicillin (100 U/ml)/streptomycin (100 μg/ml) (Gibco) in a humidified incubator at 37°C with 5% CO_2_. Neuroblastoma patient-derived NB-1691 cell line was established at St. Jude Children's Research Hospital and maintained in RPMI medium (18). All cell lines were validated by short tandem repeat (STR) using Promega PowerPlex 16 HS System. Cell lines were also regularly tested for contamination of mycoplasma by e-Myco PCR (Bulldog Bio) and were mycoplasma-free throughout this study.

### Chemicals and drugs

PGE_2_, butaprost, CAY10598, and PF04418948 were purchased from Cayman Chemical. Rolipram and forskolin were purchased from Sigma-Aldrich. Compound SID26671393 was purchased from ChemBridge. Compounds TG6-129 (SID17503974), TG4-155, and TG6-10-1 were synthesized by the Medicinal Chemistry Core at the University of Tennessee Health Science Center.

### The gene-expression database R2

Gene-expression analyses of the impact of PGE_2_ signaling-associated enzymes and receptors including two cyclooxygenases (COXs), three prostaglandin E synthases (PGESs) and four PGE_2_ receptors (EPs) on the overall survival in neuroblastoma patients were performed using a publicly available database [R2: Genomics Analysis and Visualization Platform (r2.amc.nl)]. The neuroblastoma patient datasets used in this study include Versteeg-88, Kocak-649, SEQC-498, Primary NRC-283, and Oberthuer-251.

### Correlation analysis of gene expression

For correlation analysis of multiple gene expression in human neuroblastoma samples, mRNA expression data were extracted from the neuroblastoma datasets on R2 database and analyzed using Pearson correlation coefficient.

### Cell-based cAMP assay

Cytosol cAMP was measured using a cell-based homogeneous time-resolved fluorescence resonance energy transfer (TR-FRET) method (Cisbio Bioassays) (19,20). The assay is based on the generation of a strong FRET signal upon the interaction of two fluorescent molecules: FRET donor cryptate linked with anti-cAMP antibody and FRET acceptor d2 coupled to cAMP. Endogenous cAMP synthesized by cells upon stimulation competes with fluorescence-labeled cAMP for binding to the cAMP antibody, and thus decreases the FRET signal. Human neuroblastoma cells or normal control fibroblast cells were seeded into 384-well plates with 40 μl complete medium (~4,000 cells/well) and grown overnight. The medium was completely withdrawn and 10 μl Hanks’ Balanced Salt solution (HBSS) (HyClone) supplemented with 20 μM rolipram was added into the wells to block phosphodiesterases that might metabolize cAMP. The cells were incubated at room temperature for 30 min, and then treated with vehicle or tested compounds for 5-10 min before incubation with appropriate agonists for 40 min. The cells were lysed in 10 μl lysis buffer containing the FRET acceptor cAMP-d2 and 1 min later another 10 μl lysis buffer with anti-cAMP-cryptate was added. After incubation for 1 hr at room temperature, the FRET signal was measured by a 2104 Envision Multilabel Plate Reader (PerkinElmer) with an excitation at 340/25 nm and dual emissions at 665 nm and 590 nm for d2 and cryptate (100 μs delay), respectively. The FRET signal was expressed as F665/F590×10^4^.

### Animal procedures

Athymic nude mice (4-6 weeks, female) from Charles River Laboratories were housed under a 12-hr light/dark cycle with food and water *ad libitum*. Every effort was made to lessen animal suffering. All animal procedures followed the institutional and IACUC guidelines at the University of Tennessee Health Science Center. To create flank tumors, mice were subcutaneously injected with human SK-N-AS neuroblastoma cells harboring 11q-deletion mixed with Matrigel while the animals were anaesthetized with isoflurane to enable an accurate site of injection. Tumor growth was monitored by measuring tumor volume daily using the formula: V = (width)^2^ × (length) × 0.5. After solid tumors appeared, mice were randomized and treated with either vehicle or EP2 antagonists as indicated in each experiment.

### Immunohistochemistry

Subcutaneous tumors were harvested and sectioned (7 μm) using an HM525 NX Cryostat (Thermo Scientific). The tumor tissue sections were fixed by 4% paraformaldehyde (PFA) at room temperature for 20 min and permeabilized with 0.25% Triton X-100 at room temperature for 15 min. After blocking in 10% goat serum in PBS at room temperature for 1 hr, the sections were incubated in primary antibodies at 4°C overnight: rabbit anti-Ki-67 (1:200, Biocare Medical, Cat #CRM 325B); and mouse anti-cluster of differentiation 31 (CD31) (1:200, eBioscience, Cat #11-0311-82). The sections were washed with PBS and incubated with goat anti-rabbit secondary antibody Alexa Fluor 488 (Invitrogen, Cat # A-11034) at room temperature for 2 hr and DAPI (1 μg/ml in PBS) for 10 min. Stained sections were mounted on slides using DPX Mountant (Electron Microscopy Sciences). Images were obtained using a fluorescence microscope BZ-X800 (Keyence). The fluorescence intensity was quantified using ImageJ software developed by the National Institutes of Health (NIH).

### Statistical analysis

The PGE_2_ dose-response curves were generated and EC_50_s were calculated using OriginPro software (OriginLab). Statistical analyses were performed using Prism 7.05 (GraphPad Software) by one-way/two-way ANOVA with *post-hoc* Bonferroni’s or Dunnett’s test for multiple comparisons, Fisher’s exact test, or *t*-test as indicated in each experiment. The correlation analyses were performed using Pearson correlation coefficient. *P* < 0.05 was considered statistically significant. Data are presented as mean + or ± SEM.

## RESULTS

### COX/PGES/EP signaling pathways in human neuroblastoma

To study the PGE_2_ signaling pathways in neuroblastoma, we first analyzed the gene expression data in human neuroblastoma samples from the R2 database using Kaplan-Meier estimator with *post-hoc* log-rank test. We began with the Versteeg cohort (*N* = 88) because it was the first neuroblastoma dataset publicly available on R2 platform and has been widely used. Among all nine PGE_2_ signaling-associated genes, the expression of COX-1, mPGES-1, EP1, and EP2 highly correlated with poor survival in neuroblastoma patients (*P* = 2.6E-6, 0.015, 0.016, and 6.2E-8, respectively) (**Fig. 1A and Table 1**). In line with this, survival patients had significantly lower expression of COX-1 and EP2 in their tumors than nonsurvival patients (*P* < 0.01 for COX-1; *P* < 0.001 for EP2) (**Fig. 1B**). Furthermore, all patients with tumors expressing high levels of EP2 receptor deceased in less than 30 months, whereas over 70% of patients with tumor expressing low levels of EP2 eventually survived (**Fig. 1A**). Nonsurvival patients had approximately 61% more COX-1 and 79% more EP2 expression in their tumors than survival patients (**Fig. 1B**). On the contrary, the expression of Gα_i_-coupled EP3 showed a significant positive correlation with the probability of survival in neuroblastoma patients (*P* = 1.8E-4) (**Fig. 1A**), as nonsurvival patients had about 35% less EP3 expression in their tumors compared to survival patients (*P* < 0.05) (**Fig. 1B**). Although the expression of EP1 receptor also correlated with decreased patient survival (**Fig. 1A**), overall there was no statistically significant difference in EP1 expression between survival and nonsurvival patients (**Fig. 1B**). Interestingly, both EP2 and EP4 receptors but not EP1 showed expression correlation with COX-1 in neuroblastoma tumors (*P* = 0.17663, 9.65E-7 and 3.17E-4 for EP1, EP2 and EP4, respectively). In contrast, the expression of EP3 receptor showed a trend of negative correlation with that of COX-1 (**Fig. 1C**). These interesting results together suggest that the PGE_2_ receptor EP2 is positively – whereas EP3 is negatively – associated with the increased aggressiveness of neuroblastoma.

**Table 1.** Relationship of the survival probability and expression of PGE_2_-related genes in human neuroblastoma. The relationship between survival probability of neuroblastoma and the expression of nine PGE_2_ signaling-associated genes from five major patient cohorts on R2 database platform was examined by Kaplan-Meier estimator with *post-hoc* log-rank test. Negative relationships (*P* < 0.05) are highlighted in red and positive relationships (*P* < 0.05) are in blue, with *X* and *P* values indicated. The relationship is not considered significant if *P* > 0.05. N/A: gene expression data are not available. Note that only the expression of COX-1 and EP2 had significant negative correlation with the patient survival across all neuroblastoma datasets.

| Gene | Versteeg<br>(N = 88) | Kocak<br>(N = 649) | SEQC<br>(N = 498) | NRC<br>(N = 283) | Oberthuer<br>(N = 251) |
| --- | --- | --- | --- | --- | --- |
| COX-1<br>(PTGS1) | *X* = 22.10<br>*P* = 2.6E-6 | *X* = 15.74<br>*P* = 7.2E-5 | *X* = 8.91<br>*P* = 2.8E-3 | *X* = 9.58<br>*P* = 2.0E-3 | *X* = 9.81<br>*P* = 1.7E-3 |
| COX-2<br>(PTGS2) | *X* = 2.18<br>*P* = 0.140 | *X* = 11.42<br>*P* = 7.3E-4 | *X* = 12.01<br>*P* = 5.3E-4 | *X* = 2.89<br>*P* = 0.089 | *X* = 5.64<br>*P* = 0.018 |
| mPGES-1<br>(PTGES) | *X* = 5.88<br>*P* = 0.015 | *X* = 3.38<br>*P* = 0.066 | *X* = 4.06<br>*P* = 0.044 | *X* = 3.83<br>*P* = 0.050 | N/A |
| mPGES-2<br>(PTGES2) | *X* = 2.16<br>*P* = 0.141 | *X* = 27.13<br>*P* = 1.9E-7 | *X* = 17.40<br>*P* = 3.0E-5 | *X* = 14.68<br>*P* = 1.3E-4 | *X* = 11.99<br>*P* = 5.3E-4 |
| cPGES<br>(PTGES3) | *X* = 1.35<br>*P* = 0.245 | *X* = 5.40<br>*P* = 0.020 | *X* = 41.21<br>*P* = 1.4E-10 | N/A | *X* = 36.30<br>*P* = 1.7E-9 |
| EP1<br>(PTGER1) | *X* = 5.82<br>*P* = 0.016 | *X* = 7.96<br>*P* = 4.8E-3 | *X* = 5.83<br>*P* = 0.016 | *X* = 11.63<br>*P* = 6.5E-4 | N/A |
| EP2<br>(PTGER2) | *X* = 29.31<br>*P* = 6.2E-8 | *X* = 74.19<br>*P* = 7.1E-18 | *X* = 71.56<br>*P* = 2.7E-17 | *X* = 30.78<br>*P* = 2.9E-8 | N/A |
| EP3<br>(PTGER3) | *X* = 14.05<br>*P* = 1.8E-4 | *X* = 15.96<br>*P* = 6.5E-5 | *X* = 42.00<br>*P* = 9.1E-11 | *X* = 3.70<br>*P* = 0.054 | *X* = 18.50<br>*P* = 1.7E-5 |
| EP4<br>(PTGER4) | *X* = 3.91<br>*P* = 0.048 | *X* = 23.76<br>*P* = 1.1E-6 | *X* = 23.91<br>*P* = 1.0E-6 | *X* = 7.34<br>*P* = 6.7E-3 | *X* = 8.08<br>*P* = 4.5E-3 |

**Figure 1.**
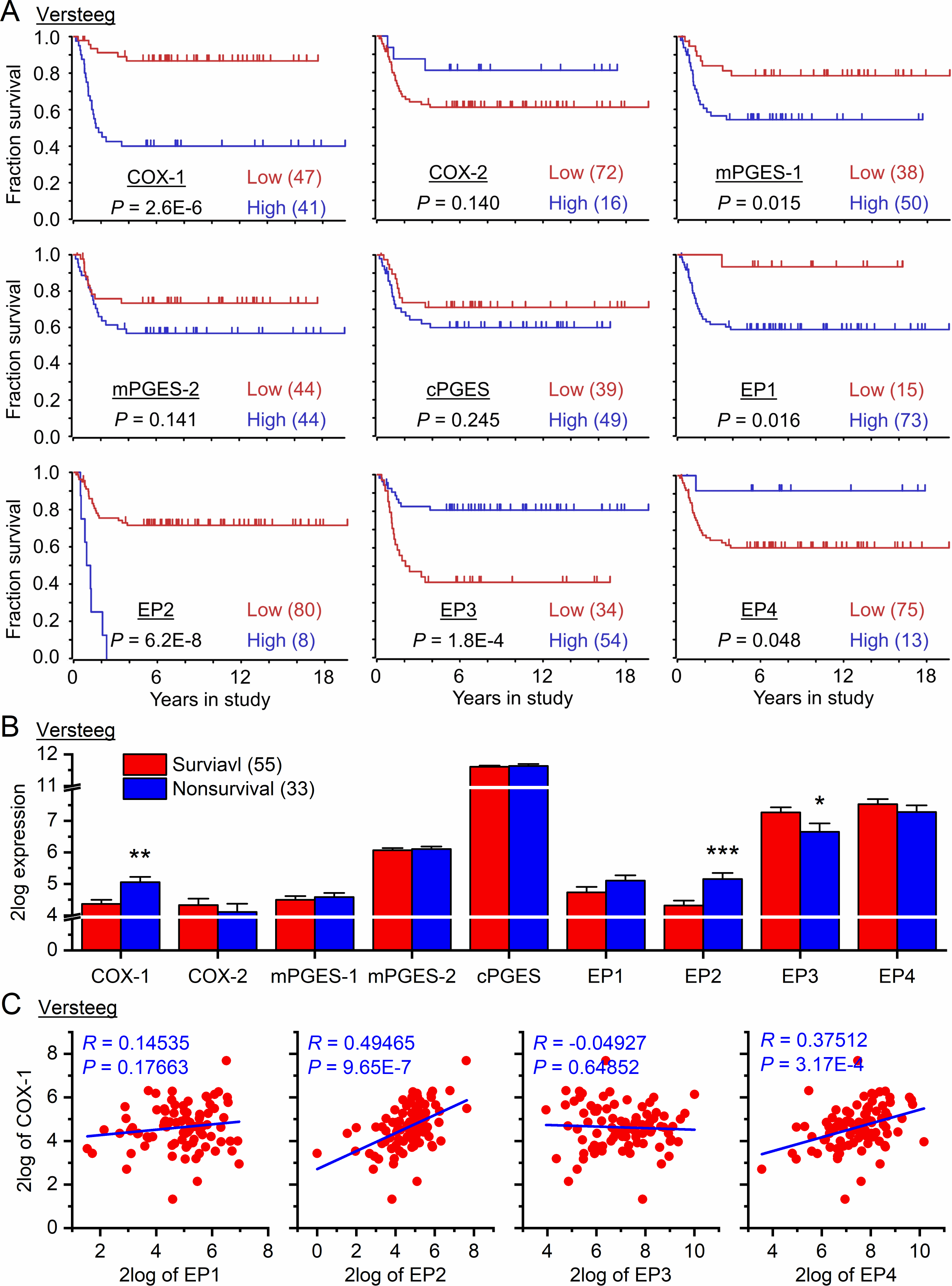
COX-1/PGE_2_/EP2 signaling axis is highly associated with human neuroblastoma. (**A**) The relationship between survival probability of neuroblastoma patients from Versteeg-88 dataset (*N* = 88) and the expression of PGE_2_ signaling-associated genes including COX-1 (*PTGS1*), COX-2 (*PTGS2*), mPGES-1 (*PTGES*), mPGES-2 (*PTGES2*), cPGES (*PTGES3*), EP1 (*PTGER1*), EP2 (*PTGER2*), EP3 (*PTGER3*), and EP4 (*PTGER4*) in their tumors was shown by Kaplan-Meier estimator with *post-hoc* log-rank test. (**B**) The expression of PGE_2_ signaling-associated genes in human neuroblastoma tissues (*N* = 55 for survival and 33 for nonsurvival patients, \**P* < 0.05; \*\**P* < 0.01; \*\*\**P* < 0.001, two-tailed unpaired *t*-test). (**C**) Pearson’s correlation coefficient analysis was performed to show the relationship of expression levels of COX-1 and EP1-EP4 receptors in human neuroblastoma tissues (*N* = 88). The gene expression and survival analyses were based on the Versteeg-88 dataset of the R2 platform.

We next examined another four large neuroblastoma cohorts (*N* > 250) on R2 platform: Kocak-649, SEQC-498, Primary-NRC-283, and Oberthuer-251 using Kaplan-Meier estimator with *post-hoc* log-rank test (**Table 1**). Consistently, we found that only the expression levels of COX-1 enzyme and EP2 receptor significantly and negatively correlated with the neuroblastoma patient survival across all cohort studies (*P* ≤ 2.8E-3 for COX-1; *P* ≤ 6.2E-8 for EP2). Similarly, mPGES-2 and cPGES mainly showed negative correlation with the patient survival in most of these neuroblastoma cohorts (*P* ≤ 5.3E-4 for mPGES-2; *P* ≤ 0.020 for cPGES). In contrast, increased expression of EP3 and EP4 receptors significantly related to the increased patient survival in the majority of these patient datasets (*P* ≤ 1.8E-4 for EP3 and *P* ≤ 0.048 for EP4). However, COX-2, mPGES-1, and EP1 showed fluctuated correlation (negative vs. positive) with the patient survival in these five large neuroblastoma cohorts (**Table 1**). Therefore, these correlation results from the five major neuroblastoma cohort studies reinforce the notion that the COX/PGES/EP2 signaling axis overall is significantly associated with the aggressiveness of human neuroblastoma.

### EP2 receptor signaling correlates with neuroblastoma-promoting factors

The activation of EP2 receptor can induce a myriad of tumor-promoting cytokines, chemokines, growth factors, and responding receptors, thereby amplifying inflammation, strengthening the tumor microenvironment, and promoting the growth of tumors in skin, colon, prostate, and brain (14,16,21–24). We next examined the expression relationships between EP2 receptor and a number of well-known tumor-associated factors via analyzing the Versteeg-88 dataset (**Table 2**). Indeed, there were significant positive correlations between EP2 receptor and the majority of examined pro-tumor mediators (29 out of 36) that might regulate tumor proliferation, survival, migration, invasion, angiogenesis and immune evasion in human neuroblastoma. These neuroblastoma-promoting factors include ALK (25), BDNF (26), CCL2/CCR2 (27), CSF-1/CSF-1R (28), CX3CL1/CX3CR1 (29), CXCL2/CXCR2 (30), CXCL12/CXCR4/CXCR7 (31), EGFR (32), IL-1β/IL1R (33), IL-6/IL6R (34,35), MIC-1 (36), MMPs (37,38), PDGFRB (13), PECAM-1 (39), POSTN (40,41), RARRES2/CMKLR1 (42), STAT3 (43), TGF-β1/TGFBR1 (44,45), and VEGF/VEGFR (46). Conversely, none of currently known anti-neuroblastoma factors (0 out of 5), including CSF-2, IFN-γ, IL-2, IL-27, and TNF-α (34,47,48), correlated with EP2 receptor in neuroblastoma tumors (*P* = 0.001) (**Table 2**). These findings from analyzing the neuroblastoma patient datasets of R2 database platform together reveal that the elevated EP2 receptor expression is highly associated with the increased malignancy of neuroblastoma tumors, leading us to hypothesize that PGE_2_ signaling via EP2 receptor might contribute to COX activity-mediated neuroblastoma growth.

**Table 2.** Expression correlation between EP2 receptor and pro- or anti-tumor factors in neuroblastoma Pro-neuroblastoma cytokines and growth factors. The expression relationship between prostaglandin EP2 receptor and a number of currently known neuroblastoma-promoting or inhibiting cytokines, chemokines, growth factors and corresponding receptors in human neuroblastoma from the Versteeg-88 dataset was examined by Pearson’s correlation coefficient analysis (*N* = 88). Twenty-nine out of these 36 pro-neuroblastoma factors show significantly positive correlations (*P* < 0.05) with EP2 receptor, which are highlighted in red, whereas none of the five anti-neuroblastoma factors correlates with EP2 (*P* = 0.001, Fisher’s exact test). Abbreviations: ALK = anaplastic lymphoma kinase; BDNF = brain-derived neurotrophic factor; CAF = cancer-associated fibroblast; CCL2 = chemokine (C-C motif) ligand 2; CCR2 = C-C chemokine receptor type 2; CMKLR1 = chemokine like receptor 1 or ChemR23 = chemerin receptor 23; CSF1 = colony stimulating factor 1; CSF1R = colony stimulating factor 1 receptor; CSF2 = colony stimulating factor 2 or granulocyte-macrophage colony-stimulating factor (GM-CSF); CX3CL1 = chemokine (C-X3-C motif) ligand 1; CX3CR1 = CX3C chemokine receptor 1; CXCL2 = chemokine (C-X-C motif) ligand 2; CXCL12 = chemokine (C-X-C motif) ligand 2; CXCR2 = C-X-C chemokine receptor type 2; CXCL12 = C-X-C motif chemokine ligand 12; CXCR4 = chemokine (C-X-C motif) receptor type 4; CXCR7 = C-X-C chemokine receptor type 7; EGFR = epidermal growth factor receptor; IL1B = interleukin 1β; IL1R1 = interleukin 1 receptor type I; IL1R2 = interleukin 1 receptor type 2; IL2 = interleukin 2; IL6 = interleukin 6; IL6R = interleukin 6 receptor; IL27 = interleukin 27; *IFNG* = interferon γ; MIC1 = macrophage inhibitory cytokine-1; MIF = macrophage migration inhibitory factor; MMP1 = matrix metalloproteinase-1; MMP2 = matrix metalloproteinase-2; MMP9 = matrix metalloproteinase-9; NK cells = natural killer cells; PDGFRB = platelet-derived growth factor receptor β; PECAM1 = platelet endothelial cell adhesion molecule-1 or cluster of differentiation 31 (CD31); POSTN = periostin or osteoblast-specific factor-2 (OSF-2); RARRES2 = retinoic acid receptor responder protein 2 or chemerin; STAT3 = signal transducer and activator of transcription 3; TGFB1 = transforming growth factor β1; TGFBR1 = transforming growth factor β receptor 1; TNF = tumor necrosis factor α; VEGF = vascular endothelial growth factor; VEGFR1 = vascular endothelial growth factor receptor 1.

| **Gene** | **R** | **P** | **Function and Mechanism in Neuroblastoma** | **Reference** |
| --- | --- | --- | --- | --- |
| *ALK* | 0.214 | 0.045 | Co-amplification with *MYCN* | Trigg and Turner, 2018 |
| *BDNF* | 0.203 | 0.058 | Survival and chemoprotection of neuroblastoma cells | Middlemas et al., 1999 |
| *CCL2* | 0.478 | 2.5E-6 | NK cells infiltration | Metelitsa et al., 2004 |
| *CCR2* | 0.432 | 2.6E-5 |  |  |
| *CSF1* | 0.401 | 1.1E-4 | Tumor growth via recruiting monocytes and macrophages | Webb et al., 2018 |
| *CSF1R* | 0.480 | 2.2E-6 |  |  |
| *CX3CL1* | 0.386 | 2.1E-4 | Bone-marrow metastasis of neuroblastoma | Nevo et al., 2009 |
| *CX3CR1* | 0.399 | 1.2E-4 |  |  |
| *CXCL2* | 0.060 | 0.579 | Tumor invasiveness | Hashimoto et al., 2009 |
| *CXCR2* | 0.309 | 3.4E-3 |  |  |
| *CXCL12* | 0.401 | 1.1E-4 | Tumor cell growth, survival and angiogenesis | Liberman et al., 2012 |
| *CXCR4* | 0.252 | 0.018 |  |  |
| *CXCR7* | 0.277 | 8.9E-3 |  |  |
| *EGFR* | 0.240 | 0.025 | Neuroblastoma cell proliferation | Ho et al., 2005 |
| *IL1B* | 0.276 | 9.2E-3 | Tumor growth and metastasis | Elaraj et al., 2006 |
| *IL1R1* | 0.539 | 6.1E-8 |  |  |
| *IL1R2* | 0.110 | 0.309 |  |  |
| *IL6* | 0.349 | 8.6E-4 | Tumor growth and metastasis; drug resistance | Pistoia et al., 2011; Ara et al., 2013; |
| *IL6R* | 0.270 | 0.011 |  |  |
| *MIC1* | 0.309 | 3.4E-3 | Regulation of the stem cell-like phenotype in neuroblastoma | Craig et al., 2016 |
| *MIF* | -0.120 | 0.266 | Upregulation of *MYCN* and angiogenic factors | Ren et al., 2004 |
| *MMP1* | 0.285 | 7.0E-3 | Tumor progression | Nyalendo et al., 2009 |
| *MMP2* | 0.437 | 2.1E-5 | Tumor metastasis | Sugiura et al., 1998 |
| *MMP9* | 0.326 | 1.9E-3 | Tumor metastasis | Sugiura et al., 1998 |
| *PDGFRB* | 0.384 | 2.2E-4 | CAF marker | Kock et al., 2018 |
| *PECAM1* | 0.469 | 4.1E-6 | Angiogenesis | Pezzolo et al., 2007 |
| *POSTN* | 0.237 | 0.026 | Tumor development | Sasaki et al., 2002;<br>Morra and Moch,<br>2011 |
| *RARRES2* | 0.350 | 8.2E-4 | Chemotaxis, cell adhesion, survival and proliferation | Tummler et al.,<br>2017 |
| *CMKLR1* | 0.321 | 2.3E-3 |  |  |
| *STAT3* | 0.239 | 0.025 | Drug resistance | Hadjidaniel et al.,<br>2017 |
| *TGFB1* | 0.438 | 2.0E-5 | Tumor escape from NK cells-mediated immune surveillance | Tran et al., 2017;<br>Castriconi et al.,<br>2013; |
| *TGFBR1* | -0.193 | 0.071 |  |  |
| *VEGFA* | 0.042 | 0.695 | Angiogenesis | Jakovljevic et al.,<br>2009 |
| *VEGFB* | 0.074 | 0.495 |  |  |
| *VEGFC* | 0.392 | 1.6E-4 |  |  |
| *VEGFR1* | 0.284 | 7.4E-3 |  |  |

Anti-neuroblastoma cytokines and growth factors
| Gene | R | P | Function and Mechanism in Neuroblastoma | Reference |
| --- | --- | --- | --- | --- |
| *CSF2* | -0.010 | 0.923 | Potentialiation of the activity of anti-neuroblastoma antibodies | Pistoia et al., 2011 |
| *IFNG* | -0.098 | 0.361 | Tumor apoptosis and inhibiting angiogenesis | Pistoia et al., 2011 |
| *IL2* | 0.123 | 0.253 | Potentialiation of the activity of anti-neuroblastoma antibodies | Salcedo et al.,<br>2009 |
| *IL27* | 0.146 | 0.175 | Induction of tumor regression | Salcedo et al.,<br>2009 |
| *TNF* | 0.134 | 0.215 | Apoptosis in neuroblastoma cells | Alvarez et al., 2011 |

### PGE_2_ mediates Gα_s_-dependent signaling via EP2 in human neuroblastoma cells

The expression of PGE_2_ signaling-associated genes including COXs, PGESs, and EPs was previously demonstrated in human neuroblastoma cells and tissues (13,17). However, gene expression does not necessarily correlate with signaling activity. To further investigate the PGE_2_/EPs/cAMP signaling in high-risk neuroblastoma, we treated human neuroblastoma cell lines SK-N-SH, SK-N-BE(2) (*MYCN*-amplification), BE(2)-C (*MYCN*-amplification), SH-SY5Y, SK-N-AS (11q-deletion), and human patient-derived NB1691 (*MYCN*-amplification) with a high concentration (10 μM) of PGE_2_, EP2 selective agonist butaprost, or EP4 selective agonist CAY10598, aiming to fully activate EP2 or EP4 receptor. The human fibroblast cell line HS68 was included in the experiment as normal control cells (18). The cytosol cAMP levels were measured by a time-resolved fluorescence energy transfer (TR-FRET) assay, in which a reduction of FRET signal indicates an increase in cAMP level. We found that PGE_2_ and butaprost, but not CAY10598, induced cAMP accumulation in all examined neuroblastoma cell lines to a degree similar to that of 100 μM forskolin (*P* < 0.001) (**Fig. 2A**), a direct activator of the adenylyl cyclase commonly used to indicate the maximal capability of the cells to produce cAMP. In contrast, stimulation with EP4 agonist CAY10598 led to more cAMP production than EP2 agonist butaprost and PGE_2_ in the normal human fibroblast cell line HS68 (**Fig. 2B**). It appears that the EP2 receptor, when compared to the much less active EP4, is a dominant Gα_s_-coupled receptor that mediates PGE_2_-initaited cAMP signaling across all these tested neuroblastoma cells regardless of the MYCN status (*P* = 0.002) (**Fig. 2C**). However, EP4 is the leading PGE_2_ receptor subtype that transduces the cAMP signaling in the normal fibroblast cells (**Fig. 2B and 2C**).

**Figure 2.**
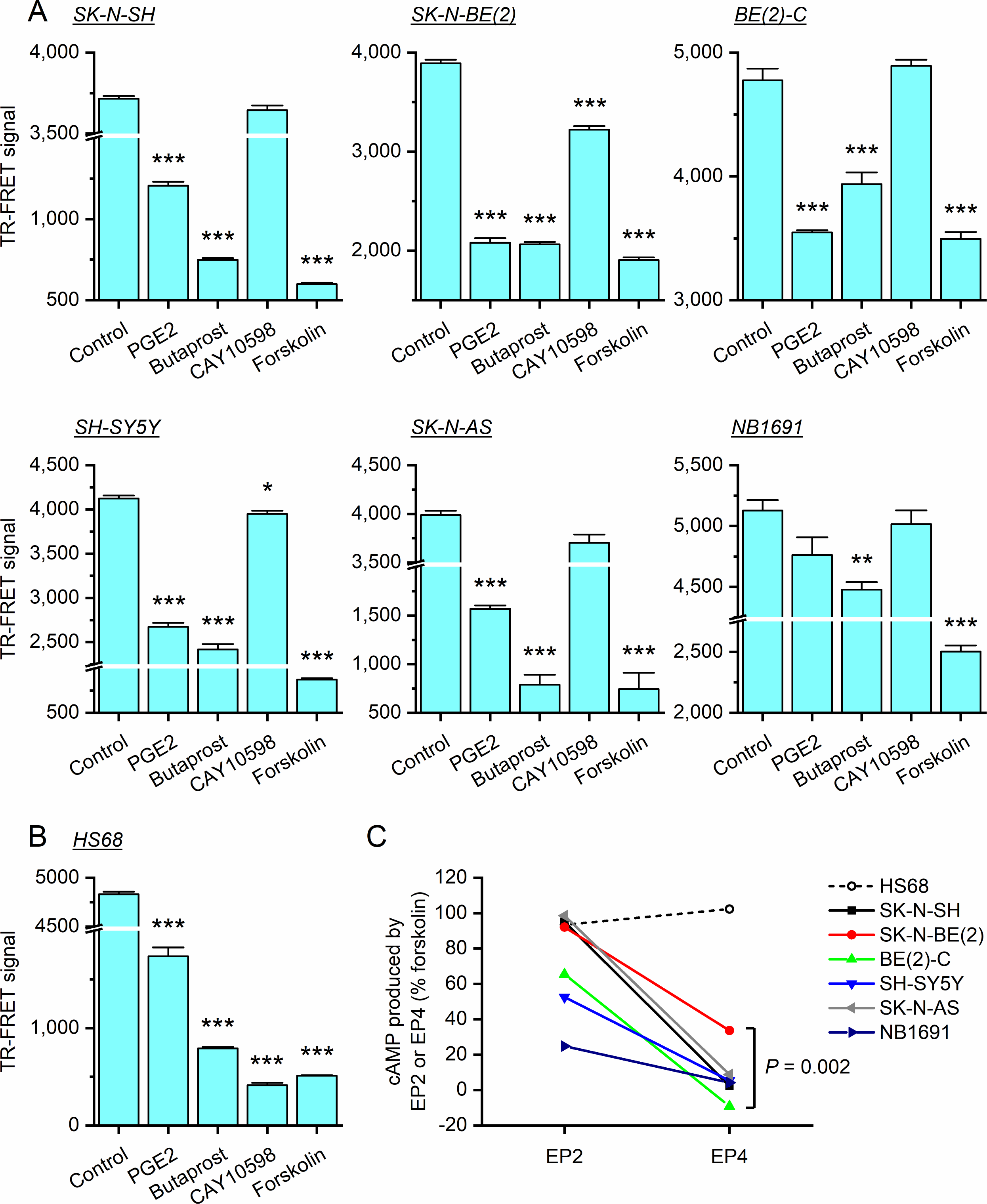
PGE_2_ mediates cAMP signaling in human neuroblastoma cells. (**A**) The human neuroblastoma cell lines SK-N-SH, SK-N-BE(2), BE(2)-C, SH-SY5Y, SK-N-AS, and patient-derived NB1691 neuroblastoma cells, and (**B**) the normal human fibroblast cell line HS68 were stimulated with PGE_2_ (10 μM), butaprost (10 μM), CAY10598 (10 μM), or forskolin (100 μM). The biosynthesis of cAMP in these cells was detected by a cell-based time-resolved fluorescence energy transfer (TR-FRET) method (*N* = 4 or 6, \**P* < 0.05; \*\**P* < 0.01; \*\*\**P* < 0.001 compared with the control group, one-way ANOVA and *post-hoc* Dunnett’s test). Data are shown as mean + SEM. (**C**) The cAMP production via EP2 receptor activation by butaprost and EP4 activation by CAY10598 in the six neuroblastoma cell lines was normalized as percent maximum response by forskolin (as 100%) and compared (*P* = 0.002, two-tailed paired *t*-test).

### Competitive antagonism on EP2/cAMP signaling in human neuroblastoma cells

We next focused on the SK-N-AS because this cell line has high *c-MYC* expression and contains 11q-deletion (49,50), a high-risk factor commonly found in human neuroblastoma. TR-FRET results using SK-N-AS cells further revealed that PGE_2_ and butaprost, but not CAY10598, activated EP2 receptor in a concentration-dependent manner and 1 μM agonists nearly maximized the cell response (**Fig. 3A**). We previously identified a small-molecule compound TG4-155 that is among the first-generation EP2-selective antagonists with high potency and selectivity (51,52). Here, TG4-155 and its analog TG6-10-1 decreased 1 μM PGE_2_-induced cAMP synthesis in SK-N-AS cells in a concentration dependent manner (**Fig. 3B**). We previously also reported another group of EP2 antagonists that have a distinct chemical scaffold and are exemplified by SID17503974 (TG6-129) and SID26671393 (53). Likewise, these two newer compounds and PF-04418948, a selective EP2 antagonist reported by Pfizer (51), showed robust inhibition on cAMP production in SK-N-AS cells stimulated by 1 μM PGE_2_. In contrast, GW627368X, an EP4-selective small-molecule antagonist had no effect on PGE_2_-induced cAMP signaling in these cells (**Fig. 3B**). All tested ccompounds inhibited PGE_2_-induced human EP2 receptor activation in SK-N-AS cells in a concentration-dependent manner, exemplified by TG6-129 and SID26671393 (**Fig. 3C**). By comparison at the same concentration (1 μM), TG4-155 showed the highest potency of EP2 antagonism in neuroblastoma cells, followed by TG6-129, PF-04418948, TG6-10-1, and SID26671393 in order, demonstrated by the rightward shift of their PGE_2_ dose-response curves compared to the control (**Fig. 3D**). Schild regression analysis revealed that our compounds TG4-155, TG6-129, TG6-10-1, and SID26671393 inhibited the EP2 receptor via a competitive mechanism and their *K*_B_ values for EP2 receptor in neuroblastoma SK-N-AS cells were 2.25 nM, 5.96 nM, 10.5 nM, and 30.3 nM, respectively (**Fig. 3E**). These TR-FRET results together revealed a robust PGE_2_/EP2/cAMP signaling that ubiquitously exists in human neuroblastoma cells.

**Figure 3.**
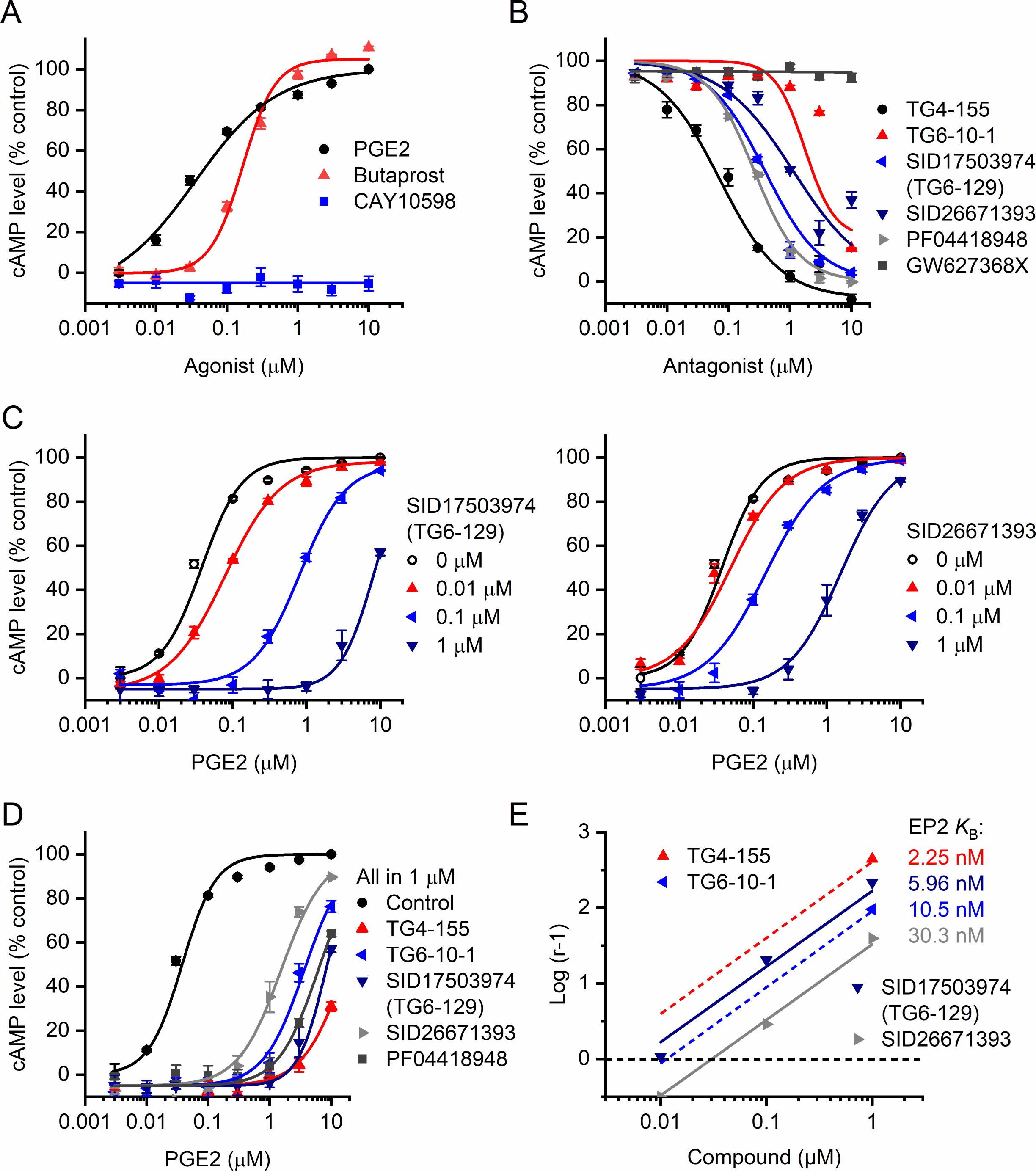
EP2 receptor is the major mediator of PGE_2_/cAMP signaling in human neuroblastoma cells. (**A**) PGE_2_ and EP2 agonist butaprost, but not EP4 agonist CAY10598, induced cAMP synthesis in human neuroblastoma cells SK-N-AS in a concentration-dependent manner. The calculated PGE_2_ EC_50_ was 0.04 μM; butaprost EC_50_ was 0.17 μM. (**B**) Compounds TG4-155, TG6-10-1, TG6-129 (SID17503974), SID26671393, and PF04418948, but not EP4 antagonist GW627368X, inhibited 1 μM PGE_2_-induced cAMP production in human neuroblastoma cells SK-N-AS in a concentration-dependent manner. IC_50_s: 0.07 μM for TG4-155; 1.75 μM for TG6-10-1; 0.39 μM for TG6-129; 1.13 μM for SID26671393; 0.26 μM for PF04418948. (**C**) Compounds TG6-129 (**left**) and SID26671393 (**right**) inhibited PGE_2_-induced human EP2 receptor activation in SK-N-AS cells in a concentration-dependent manner. PGE_2_ EC_50_s: 0.04 μM for control; 0.08, 0.81 and 8.22 μM for 0.01, 0.1 and 1 μM TG6-129, respectively; 0.05, 0.15 and 1.54 μM for 0.01, 0.1 and 1 μM SID26671393, respectively. (**D**) EP2 antagonists TG4-155, TG6-10-1, TG6-129, SID26671393, and PF04418948 (all 1 μM) showed robust inhibition on PGE_2_-induced cAMP production in SK-N-AS cells. PGE_2_ EC_50_s: 0.04 μM for control; 16.81 μM for TG4-155; 3.65 μM for TG6-10-1; 8.22 μM for TG6-129; 1.54 μM for SID26671393; 6.17 μM for PF04418948. (**E**) All tested compounds inhibited EP2 receptor in neuroblastoma cells via a competitive mechanism confirmed by Schild regression analyses. TG4-155 *K*_B_: 2.25 nM; TG6-10-1 *K*_B_: 10.5 nM; TG6-129 *K*_B_: 5.96 nM; SID26671393 *K*_B_: 30.3 nM. All data are shown as mean ± SEM (*N* = 4).

### Inhibiting EP2 receptor suppresses neuroblastoma cell growth *in vivo*

The dominant role for EP2 receptor in PGE_2_-initiated cAMP signaling in human neuroblastoma cells motivated us to test the hypothesis that EP2 receptor activation might contribute to the proliferation of neuroblastoma *in vivo*. We used SK-N-AS cells to generate subcutaneous tumors in athymic nude mice, because SK-N-AS is a 11q-deleted neuroblastoma cell line showing aggressive proliferation *in vivo* and thus has been widely used in xenograft models (9,13). We first chose compound TG6-10-1 to study the effect of pharmacological inhibition of EP2 receptor on neuroblastoma cells *in vivo* in that it has an improved *in vivo* half-life (1.6 hr) in mice than its prototype compound TG4-155 (0.6 hr) (52,54) (**Fig. 4A**). Systemic administration of TG6-10-1 was previously shown to suppress the tumor growth in glioma xenograft models (16). With oral administration in mice (10 mg/kg), the concentration of TG6-10-1 in plasma is projected to be above its EP2 Schild *K*_B_ value (4.71 ng/ml) in neuroblastoma cells for over 8 hr (**Fig. 4B**). Therefore, two doses of TG6-10-1 per day should be sufficient to achieve adequate drug exposure in this study. TG6-10-1 has high gut-blood barrier permeability revealed by QikProp (**Fig. 4C**) (Schrödinger Release 2019-4: QikProp, Schrödinger, LLC, New York, NY, 2019), indicative of the feasibility of its oral dosing. To create subcutaneous tumors, the SK-N-AS cells were injected into the flank sites of athymic nude mice. After solid tumors were developed into palpable size, animals were randomized and treated with either vehicle or compound TG6-10-1 (10 mg/kg, p.o.) twice daily. The tumor volumes were measured on a daily basis and compared. During the 13-day treatment, we found a modest decrease in the calculated size of tumors formed by SK-N-AS cells in the TG6-10-1-treated mice when compared to their vehicle-treated peers (*P* < 0.05 at day 11; *P* < 0.001 at day 13) (**Fig. 4D**).

**Figure 4.**
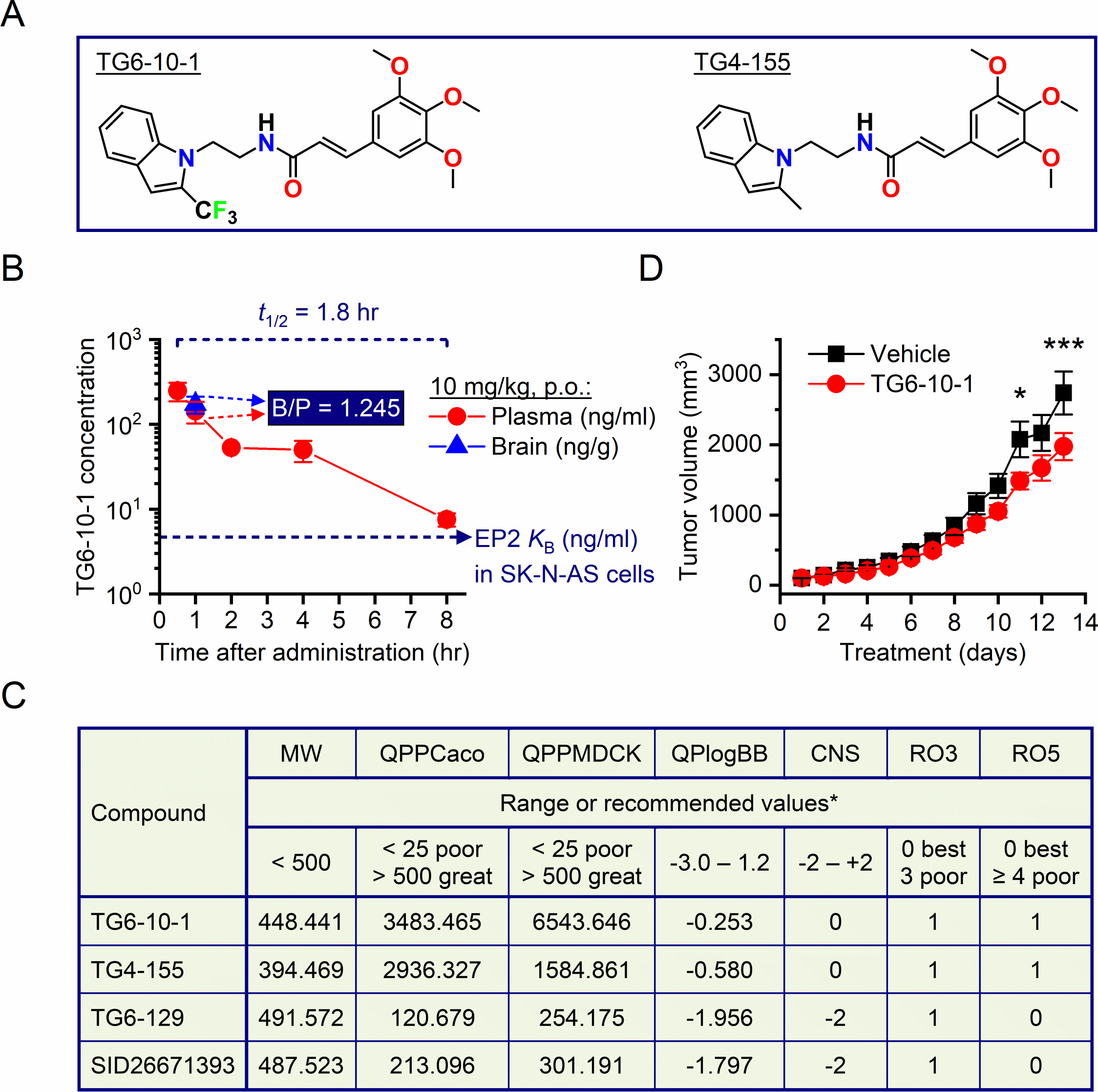
Pharmacological inhibition of EP2 receptor by TG6-10-1 slows neuroblastoma growth. (**A**) Chemical structures of EP2-selective antagonists TG6-10-1 and TG4-155. (**B**) After oral administration to mice (10 mg/ml, p.o.), compound TG6-10-1 showed a plasma half-life (*t*_1/2_) of 1.8 hr and a brain-to-plasma concentration ratio of 1.245 ± 0.107, measured 1 hr after administration (16). Data are shown as mean ± SEM (*N* = 3). The compound Schild *K*_B_ value for EP2 receptor in neuroblastoma SK-N-AS cells is also indicated: 4.71 ng/ml or 10.5 nM. (**C**) ADME properties (absorption, distribution, metabolism, excretion) of our novel EP2 antagonists were determined by QikProp. MW = molecular weight; QPPCaco = apparent Caco-2 cell permeability (nm/s) as a model for the gut-blood barrier; QPPMDCK = apparent MDCK cell permeability (nm/s) as a mimic for the blood-brain barrier; QPlogBB = brain/blood partition coefficient; CNS = central nervous system (CNS) activity on a −2 (inactive) to +2 (active) scale; RO3 = Jorgensen rule of three, an indication of oral availability; RO5 = Lipinski rule of five, an indication of drug-likeness. *Range is for 95% of known drugs. (**D**) Female athymic nude mice (4 weeks) were inoculated with human neuroblastoma cells SK-N-AS (1×10^7^ cells/site). After solid tumors were developed, vehicle (10% PEG 200 + 0.5% methylcellulose) or selective EP2 antagonist TG6-10-1 was administered (10 mg/kg, p.o, b.i.d.). Tumor growth was monitored by measuring tumor volume daily using the formula: V = (width)^2^ × (length) × 0.5 and compared [*N* = 8, F(1, 14) = 3.384, *P* = 0.087; multiple comparisons: \**P* < 0.05 at day 11 and \*\*\**P* < 0.001 at day 13, two-way ANOVA and *post-hoc* Bonferroni’s test for multiple comparisons]. Data are shown as mean ± SEM.

### Suppressing neuroblastoma by a highly bioavailable brain-impermeable EP2 antagonist

The moderate suppression on neuroblastoma growth by compound TG6-10-1 encouraged us to further test our EP2 antagonists with better pharmacokinetic (PK) and pharmacodynamic (PD) profiles. We next chose to test compound TG6-129 in neuroblastoma xenograft models for several reasons: 1) it has a completely different chemical structure than TG4-155 and TG6-10-1 that we previously used in other animal studies for CNS conditions (**Figs. 4A and 5A**) (52,54–58); 2) it has a longer terminal half-life (2.7 hr) in mice than compounds TG4-155 and TG6-10-1 (**Fig. 5B**) (52,54,59); 3) with systemic administration in mice TG6-129 has very limited presence (< 2%) in the brain (**Fig. 5B**) (53), suggesting that its effects would largely remain in the periphery. This is important for EP2-targeting therapeutics because the receptor regulates some fundamental physiological functions of the brain, such as neuroplasticity, cognition, and emotion (60,61). On the contrary, TG6-10-1 has high brain-penetration (**Fig. 4B and 4C**), indicating that the compound mainly concentrates within the brain after systemic administration. Indeed, QikProp analysis suggests that TG6-10-1 has a considerably higher CNS activity score than TG6-129 (**Fig. 4C**). Thus, higher doses of TG6-10-1 (> 10 mg/kg) may not be ideal to target peripheral EP2 receptor in neuroblastoma due to the potential risk in compromising normal brain functions.

**Figure 5.**
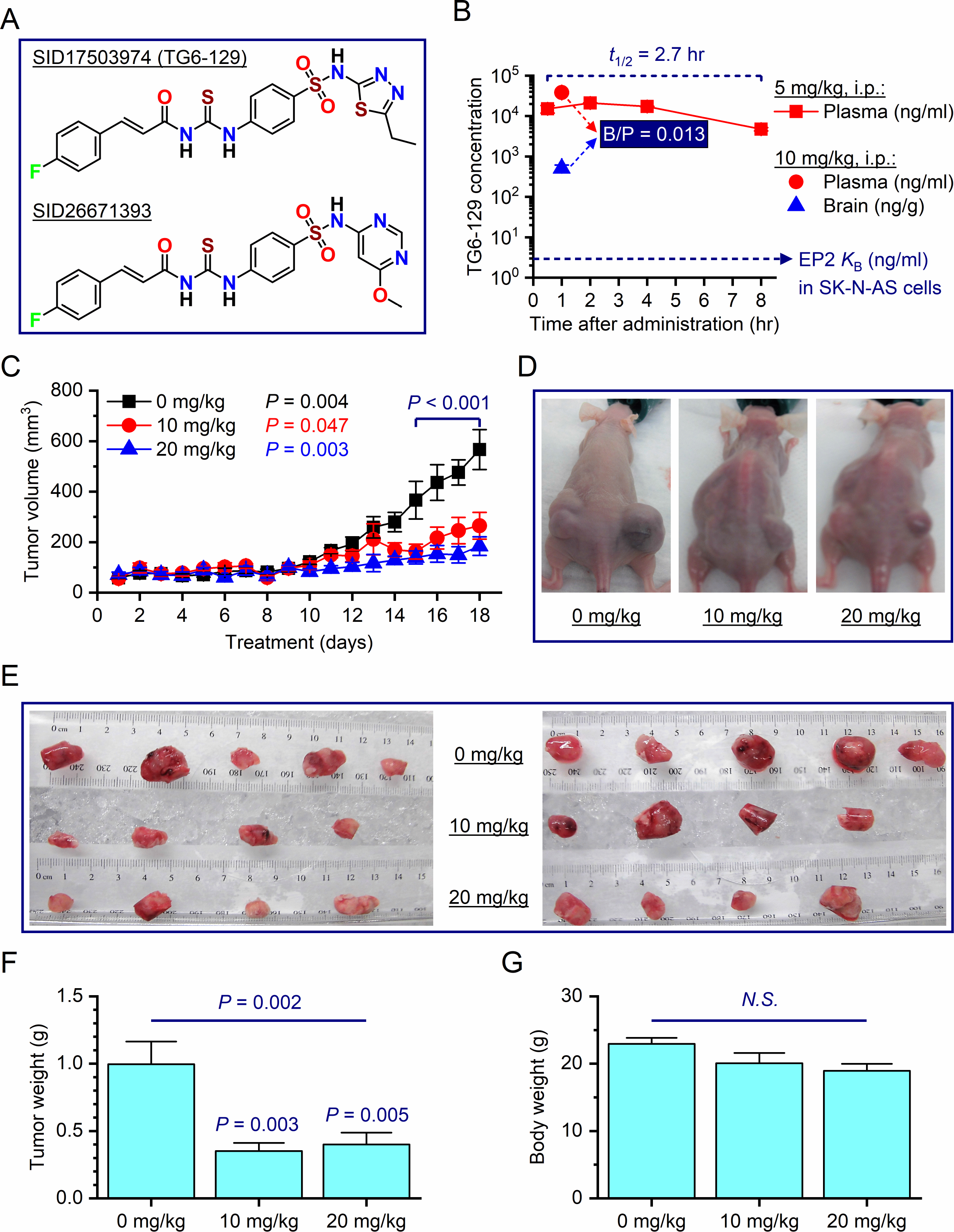
Peripheral EP2 receptor inhibition impairs neuroblastoma growth *in vivo*. (**A**) Chemical structures of EP2-selective antagonists SID17503974 (TG6-129) and SID26671393. (**B**) After systemic administration to mice (5 mg/kg, i.p.), compound TG6-129 was highly bioavailable and showed a plasma terminal half-life (*t*_1/2_) of 2.7 hr (53). One hour after intraperitoneal injection with a dose of 10 mg/kg in mice, TG6-129 showed a brain-to-plasma concentration ratio of 0.013 ± 0.002; the compound was undetectable in the brain 2 hr after injection. Data are shown as mean ± SEM (*N* = 3). The compound Schild *K*_B_ value for EP2 receptor in neuroblastoma SK-N-AS cells is also indicated: 2.93 ng/ml or 5.96 nM. (**C**) Human neuroblastoma cells SK-N-AS were inoculated into athymic nude mice (female, 6 weeks) with two injection sites per animal: 5×10^6^ cells and 10×10^6^ cells on each flank side. After solid tumors were developed, vehicle (4% DMSO + 80% PEG 400 + 16% H_2_O) or selective EP2 antagonist TG6-129 (10 or 20 mg/kg, i.p.) was administered daily for 18 consecutive days. Tumor growth was monitored by measuring tumor volume daily using the formula: V = (width)^2^ × (length) × 0.5. The SK-N-AS xenograft tumors formed by 5×10^6^ cells and 10×10^6^ cells did not significantly differ in size, so they were combined for comparisons between treatment groups [*N* = 8-10, *F* (2, 23) = 7.043, *P* = 0.004; multiple comparisons: *P* = 0.047 and 0.003 for 10 mg/kg treatment and 20 mg/kg treatment compared to control, respectively, two-way ANOVA and *post-hoc* Dunnett’s multiple comparisons test]. Data are shown as mean ± SEM. (**D**) Representative digital photographs showing tumors formed by SK-N-AS cells in mice treated with vehicle or TG6-129. (**E**) Tumors were harvested after 18-day treatment and displayed for comparisons. (**F**) All tumors were weighed and compared between treatment groups [*N* = 8-10, *F* (2, 23) = 8.645, *P* = 0.002; multiple comparisons: *P* = 0.003 for 10 mg/kg treatment and 0.005 for 20 mg/kg treatment compared to control, one-way ANOVA with *post-hoc* Dunnett’s multiple comparisons test]. Data are shown as mean + SEM. (**G**) Body weights of tumor-bearing mice before tumor harvest (*N.S.*: not significant, one-way ANOVA). Data are shown as mean + SEM.

To test TG6-129 in neuroblastoma xenografts, each mouse was inoculated with SK-N-AS cells into two different flank sites. After solid tumors were developed, animals were randomized and treated with compound TG6-129 (0, 10 or 20 mg/kg, i.p.). The mice were treated only once daily because, with a single systemic injection (10 mg/kg, i.p.) in mice, the projected concentration of TG6-129 in the circulatory system should be well above its EP2 Schild *K*_B_ value (2.93 ng/ml) in neuroblastoma cells for over 24 hr (**Fig. 5B**). Importantly, with this treatment paradigm, any undesirable effect of EP2 inhibition in the CNS would be negligible, as the compound was only detectable during the first hour after the systemic administration (**Fig. 5B**). The tumor volumes were measured on a daily basis and compared between treatment groups. After 18 days of treatment, tumors were then collected, weighed, and analyzed. Interestingly, TG6-129-treated mice (both 10 and 20 mg/kg) showed a significant decrease in tumor size when compared to vehicle-treated mice (*P* < 0.001 at days 15-18) (**Fig. 5C and 5D**). The weight of SK-N-AS tumors on average was decreased to 35% by treatment with 10 mg/kg TG6-129 (*P* = 0.003) and to 40% with treatment by 20 mg/kg TG6-129 (*P* = 0.005), when compared to vehicle treatment (**Fig. 5E and 5F**). Other than the tumor burden, all mice were overall healthy without showing any behavioral abnormality or considerable weight loss by the treatment with TG6-129 (**Fig. 5G**). Our data together suggest that PGE_2_ cAMP signaling via EP2 receptor is involved in neuroblastoma tumor growth and Gα_s_-coupled EP2 receptor might represent a novel molecular target for neuroblastoma treatment.

### Blocking EP2 reduces proliferating cells and the microvessel density in neuroblastoma

We next utilized immunohistochemistry to examine the effects of EP2 antagonist TG6-129 on the proliferative index of neuroblastoma cells *in vivo*. Immunostaining for Ki-67, a nuclear protein commonly used as a cellular marker for proliferation, was performed to identify proliferating cells in subcutaneous neuroblastoma tissues. We found that the systemic treatment with TG6-129 substantially decreased the proliferation of tumors formed by SK-N-AS cells in a dose-dependent manner: 25% reduction by 10 mg/kg (*P* = 0.004) and 55% reduction by 20 mg/kg (*P* < 0.001) (**Fig. 6A**). These findings indicated that the PGE_2_ signaling via EP2 receptor promotes the tumorigenic potential of neuroblastoma cells.

**Figure 6.**
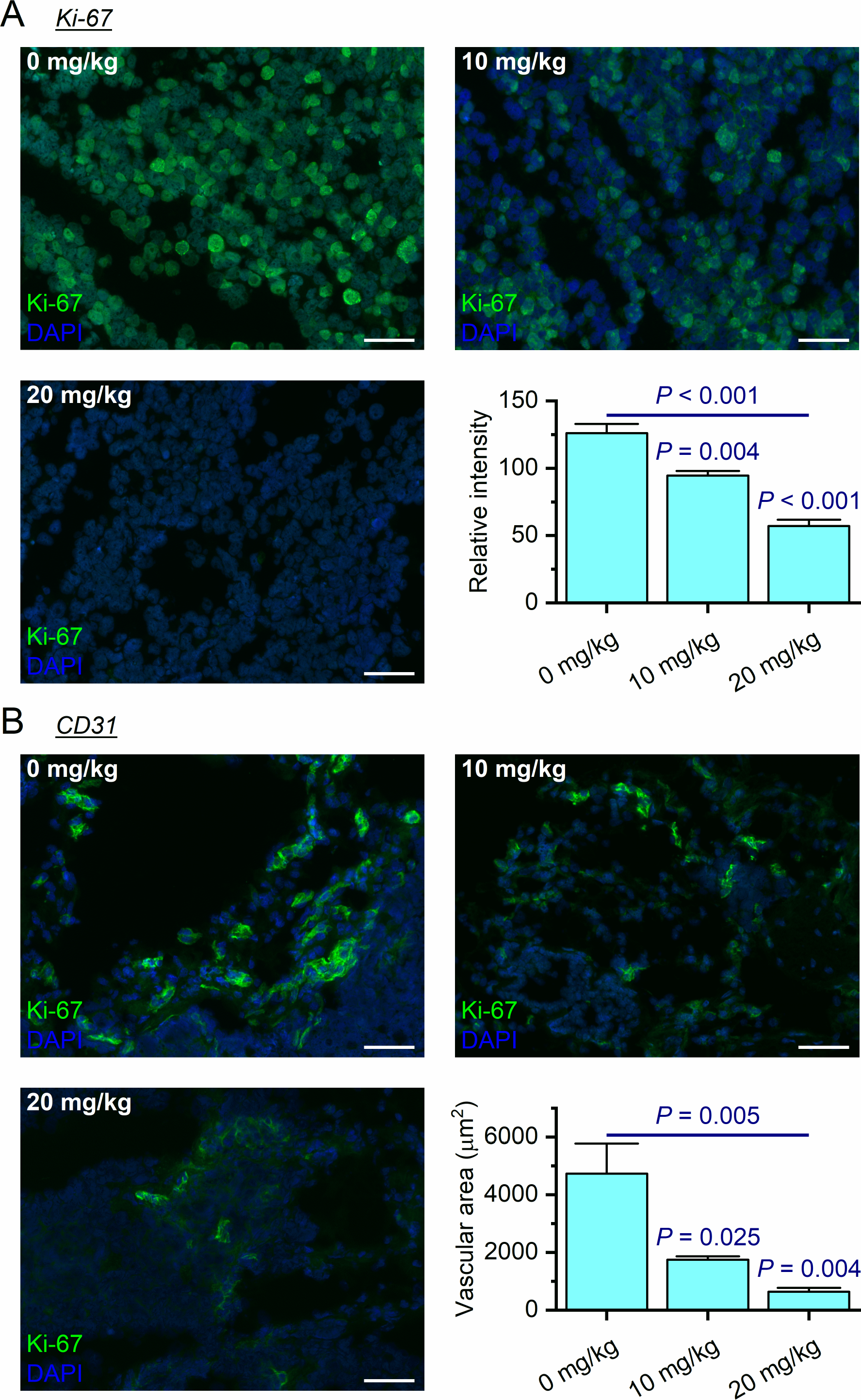
Blocking EP2 receptor decreases proliferating cells and the microvessel density. (**A**) Immunostaining for Ki-67 (green fluorescence) was performed to identify proliferating cells in subcutaneous neuroblastoma tissues. Ki-67 expression levels were measured via quantifying the fluorescence intensity using ImageJ software [*N* = 4-5, *F* (2, 10) = 39.87, *P* < 0.001; multiple comparisons: *P* = 0.004 for 10 mg/kg treatment and < 0.001 for 20 mg/kg treatment compared to control]. (**B**) Immunostaining for CD31/PECAM-1 (green fluorescence) was utilized to indicate the microvessel density in subcutaneous tumors. CD31 levels were assessed via quantifying the fluorescence intensity [*N* = 4-5, *F* (2, 10) = 9.248, *P* = 0.005; multiple comparisons: *P* = 0.025 for 10 mg/kg treatment and < 0.004 for 20 mg/kg treatment compared to control]. Note that nuclei within each tumor were counterstained with DAPI (blue fluorescence). Data are shown as mean + SEM. Scale bar = 50 μm.

Microvascular proliferation is a hallmark of higher malignancy and a cardinal characteristic of neuroblastoma especially in the advanced and aggressive stages (62). It is often indicated by the expression of the platelet endothelial cell adhesion molecule-1 (PECAM-1) or cluster of differentiation 31 (CD31), a common biomarker for angiogenesis in various tumors (63). We were thereby interested in determining whether PGE_2_ signaling via EP2 receptor contributes to the elevation of PECAM-1/CD31 in neuroblastoma. We first examined the expression relationships between PECAM-1/CD31 and the nine PGE_2_-associated genes in human neuroblastoma from the five major patient cohort studies on R2 database platform by Pearson’s correlation coefficient analysis (**Table 3**). Quite surprisingly, only the EP2 receptor consistently displayed positive correlation in expression with PECAM-1/CD31 across all neuroblastoma patient datasets (*P* ≤ 8.1E-3). To determine whether EP2 inhibition can decrease the angiogenesis in neuroblastoma, we then performed PECAM-1/CD31 immunofluorescent staining of xenograft tumor samples and quantitated their vascular areas. We found that the systemic treatment with TG6-129 largely decreased the vascular areas in SK-N-AS cells-derived tumors in a dose-dependent manner: 63% reduction by 10 mg/kg TG6-129 (*P* = 0.025) and 87% reduction by 20 mg/kg TG6-129 (*P* = 0.004) (**Fig. 6B**). Together with the substantial correlation in expression between EP2 receptor and PECAM-1/CD31 in human neuroblastoma (**Table 3**), our results revealed an important role for EP2 receptor in microvascular proliferation of high-risk neuroblastoma.

**Table 3.**
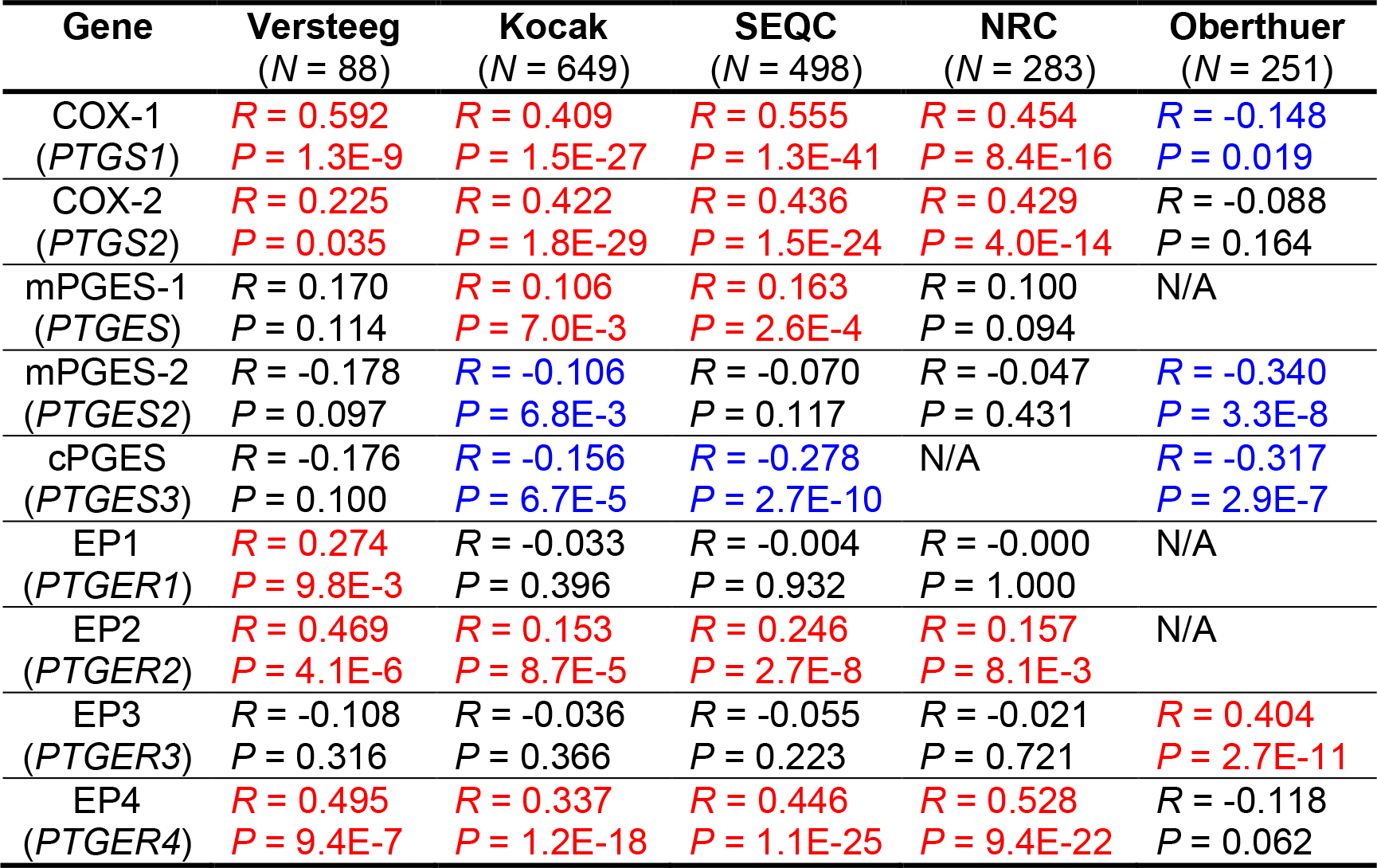
Expression correlation between COXs/PGESs/EPs and PECAM-1/CD31 in human neuroblastoma. The expression relationship between PECAM-1/CD31 and the nine PGE_2_-associated genes in human neuroblastoma from five major patient cohorts on R2 database platform was examined by Pearson’s correlation coefficient analysis. Positive correlations (*P* < 0.05) are highlighted in red and negative correlations (*P* < 0.05) are in blue. N/A: gene expression data are not available. Note that only the EP2 displayed positive correlation in expression with PECAM-1/CD31 across all neuroblastoma datasets.

## DISCUSSION

We performed a series of comprehensive analyses of gene expression and survival probability on several large cohort studies of neuroblastoma patients, which led us to hypothesize that PGE_2_ signaling via EP2 receptor plays an essential role in COX activity-associated neuroblastoma growth. We tested this hypothesis using pharmacological approaches and demonstrated that EP2 receptor is the dominant Gα_s_-coupled receptor that mediates COX/PGES/PGE_2_/cAMP signaling in both *MYCN*-amplification and 11q-deletion neuroblastoma cells. Our results from *in vivo* study further show that the EP2 receptor inhibition by a novel brain-impermeable small-molecular antagonist TG6-129 substantially decreased the aggressiveness of neuroblastoma cells and the associated angiogenesis in neuroblastoma xenografts. These findings establish PGE_2_ receptor EP2 as an appealing target for novel therapeutics aimed at suppressing the development of high-risk neuroblastoma in children.

COX activity is often elevated in tumor tissues, and its expression level is highly correlated with tumor aggressiveness (4,10). COX enzyme thus was once considered as a favorable therapeutic target for various cancers. However, the feasibility of blocking COX cascade using nonsteroidal anti-inflammatory drugs (NSAIDs) or COX-2-selective inhibitors (coxibs) to interrupt the tumor progression has been greatly challenged. Long-term use of these drugs can dramatically increase the potential risk of adversative effects particularly on gastrointestinal tract and microvascular systems that may lead to myocardial infarction, stroke, and death (64). The past decade has already witnessed a mounting recognition of these fatal consequences and the subsequent withdrawal of two legendary COX-2-targeting drugs rofecoxib and valdecoxib. These discouraging outcomes are not unanticipated, since COX activity leads to the synthesis of five types of prostanoids that in turn can activate a total of nine GPCRs, implementing a myriad of detrimental as well as beneficial actions (10,11,22,65). For instance, COX inhibition can decrease the systemic prostacyclin [also termed prostaglandin I2 (PGI_2_)], another enzymatic product of COX acts as a vasodilator and platelet inhibitor, thereby increasing cardiovascular risk (66). This monumental lesson – though disheartening – inspired us and others to seek the next-generation therapeutic targets from the COX downstream prostanoid synthases or receptors (15,67). As such, blocking mPGES-1 enzyme by a selective small-molecule inhibitor has been proposed as an alternative strategy to suppress neuroblastoma tumor growth, which is considered more specific than COX inhibition because it only disrupts PGE_2_ synthesis without affecting other COX-derived biolipids like PGI_2_ (13). Likewise, in this work, we presented evidence that targeting PGE_2_ receptor EP2 might represent another feasible therapeutic strategy for neuroblastoma. Blocking EP2 receptor activation potentially provides even higher therapeutic specificity than inhibiting mPGES-1 as it should not affect other three PGE_2_ receptor subtypes, particularly Gα_i_-coupled EP3, given that its expression is presumably correlated with the decreased aggressiveness of neuroblastoma (**Fig. 1A and 1B**).

PGE_2_ has been reported to regulate tumorigenesis through all four EP receptors, among which the Gα_s_-coupled EP2 and EP4 receptors have been mostly studied for their potential roles in the development and progression of tumors including those of breast, colon, lung, ovary, prostate, skin, and stomach (67–74). It appears that EP2 and EP4 receptors are ubiquitously present in most tumors and likely function synergistically to enhance cancer cell activities because both receptors in a very similar way can trigger cAMP signaling as well as G protein-independent pathway (15,67). However, in contrast to EP2 receptor’s tumor-promoting roles in neuroblastoma, the EP4 signaling appears to be positively associated with the probability of survival in neuroblastoma patients (**Fig. 1A and Table 1**); the EP4 receptor expression in neuroblastoma displayed a tendency to increase (19%) in survival patients when compared to nonsurvival patients (**Fig. 1B**). Indeed, the PGE_2_-promoted cAMP singling in various human neuroblastoma cell lines is mainly mediated by the EP2 receptor, although some of these cells might also express functional EP4 at relatively low levels (**Fig. 2A**). Conversely, EP4 is the dominant PGE_2_ receptor subtype for cAMP production in normal human fibroblast cells (**Fig. 2B**). Nonetheless, whether the EP4 receptor signaling down- or up-regulates the growth of neuroblastoma remains to be determined in the future.

PGE_2_ singling via EP2 receptor has emerged as an essential contributor to tumor growth engaging mechanisms including but not limited to: 1) inducing reactive mediators for tumor cell growth including pro-inflammatory cytokines and growth factors (14,22,23,75); 2) promoting angiogenesis via activating vascular endothelial growth factor (VEGF) and fibroblast growth factor (FGFR) (76–78); 3) creating immunosuppressive microenvironments that allow tumor cells to escape immunosurveillance (24,79). The molecular mechanisms by which PGE_2_/EP2 signaling promotes the progression of neuroblastoma remain elusive. However, we previously revealed that EP2 receptor activation can induce pro-inflammatory cytokines (14,52,80). Indeed, we showed here a positive correlation between EP2 receptor and a flock of essential tumor-promoting cytokines, chemokines and growth factors in human neuroblastoma tumors, such as ALK, CCL2/CCR2, CSF-1/CSF-1R, CX3CL1/CX3CR1, CXCR2, CXCL12/CXCR4/CXCR7, EGFR, IL-1β/IL1R, IL-6/IL6R, MIC-1, MMPs, PDGFRB, PECAM-1, POSTN, RARRES2/CMKLR1, STAT3, TGF-β1, and VEGFC/VEGFR1, but not the currently known anti-neuroblastoma factors including CSF-2, IFN-γ, IL-2, IL-27, and TNF-α (**Table 2**). In addition, treatment with EP2 antagonist TG6-129 decreased the PECAM-1/CD31 levels in high-risk neuroblastoma xenografts, indicative of a role of EP2 receptor in promoting microvascular proliferation of neuroblastoma. Whether the EP2 receptor activation also contributes to immunosuppressive microenvironment in neuroblastoma remains an important topic for future studies.

In summary, this pharmacological study, which was fine guided by PK and PD properties of the tested compounds, provides proof-of-concept evidence that PGE_2_ signaling via EP2 receptor might represent a novel anti-inflammatory target for the treatment of neuroblastoma with various high-risk factors. Our findings raise a notion that EP2 antagonism is an alternative strategy to inhibiting PGE_2_ synthesis for chemoprevention of childhood cancers with higher therapeutic specificity. Considering its significant therapeutic effects in high-risk neuroblastoma xenografts, the favorable PK/PD profiles, and its high drug-likeness based on the Lipinski’s rule of five (**Fig. 4C**), our current lead EP2 antagonist TG6-129 is well positioned as a prime candidate for progression to more intense pre-clinical and clinical studies.

## ACKNOWLEDGMENTS

We thank Dr. Jiawang Liu, Director of the Medicinal Chemistry Core at the University of Tennessee Health Science Center, for synthesizing compounds used in this study. This work was supported by the National Institutes of Health (NIH)/National Institute of Neurological Disorders and Stroke (NINDS) Grants R01NS100947 (to J.J.) and R21NS109687 (to J.J.), the American Cancer Society Research Scholar Grant 130421-RSG-17-071-01-TBG (to J.Y.), and the National Cancer Institute (NCI) Grants R03CA212802 (to J.Y.) and R01CA229739 (to J.Y).

## REFERENCES

1. Maris JM. Recent advances in neuroblastoma. N Engl J Med 2010;362:2202–11.

2. Caren H, Kryh H, Nethander M, Sjoberg RM, Trager C, Nilsson S, et al. High-risk neuroblastoma tumors with 11q-deletion display a poor prognostic, chromosome instability phenotype with later onset. Proceedings of the National Academy of Sciences of the United States of America 2010;107:4323–8.

3. Mlakar V, Jurkovic Mlakar S, Lopez G, Maris JM, Ansari M, Gumy-Pause F. 11q deletion in neuroblastoma: a review of biological and clinical implications. Molecular cancer 2017;16:114.

4. Mantovani A, Allavena P, Sica A, Balkwill F. Cancer-related inflammation. Nature 2008;454:436–44.

5. Grivennikov SI, Greten FR, Karin M. Immunity, inflammation, and cancer. Cell 2010;140:883–99.

6. Qiu J, Shi Z, Jiang J. Cyclooxygenase-2 in glioblastoma multiforme. Drug discovery today 2017;22:148–56.

7. Carlson LM, Kogner P. Neuroblastoma-related inflammation: May small doses of aspirin be suitable for small cancer patients? Oncoimmunology 2013;2:e24658.

8. Carlson LM, Rasmuson A, Idborg H, Segerstrom L, Jakobsson PJ, Sveinbjornsson B, et al. Low-dose aspirin delays an inflammatory tumor progression in vivo in a transgenic mouse model of neuroblastoma. Carcinogenesis 2013;34:1081–8.

9. Larsson K, Kock A, Idborg H, Arsenian Henriksson M, Martinsson T, Johnsen JI, et al. COX/mPGES-1/PGE2 pathway depicts an inflammatory-dependent high-risk neuroblastoma subset. Proceedings of the National Academy of Sciences of the United States of America 2015;112:8070–5.

10. Wang D, Dubois RN. Eicosanoids and cancer. Nature reviews Cancer 2010;10:181–93.

11. Hirata T, Narumiya S. Prostanoid receptors. Chemical reviews 2011;111:6209–30.

12. Samuelsson B, Morgenstern R, Jakobsson PJ. Membrane prostaglandin E synthase-1: a novel therapeutic target. Pharmacological reviews 2007;59:207–24.

13. Kock A, Larsson K, Bergqvist F, Eissler N, Elfman LHM, Raouf J, et al. Inhibition of Microsomal Prostaglandin E Synthase-1 in Cancer-Associated Fibroblasts Suppresses Neuroblastoma Tumor Growth. EBioMedicine 2018;32:84–92.

14. Jiang J, Dingledine R. Role of prostaglandin receptor EP2 in the regulations of cancer cell proliferation, invasion, and inflammation. J Pharmacol Exp Ther 2013;344:360–7.

15. Jiang J, Qiu J, Li Q, Shi Z. Prostaglandin E2 Signaling: Alternative Target for Glioblastoma? Trends in cancer 2017;3:75–8.

16. Qiu J, Li Q, Bell KA, Yao X, Du Y, Zhang E, et al. Small-molecule inhibition of prostaglandin E receptor 2 impairs cyclooxygenase-associated malignant glioma growth. British journal of pharmacology 2019;176:1680–99.

17. Rasmuson A, Kock A, Fuskevag OM, Kruspig B, Simon-Santamaria J, Gogvadze V, et al. Autocrine prostaglandin E2 signaling promotes tumor cell survival and proliferation in childhood neuroblastoma. PloS one 2012;7:e29331.

18. Yang J, Milasta S, Hu D, AlTahan AM, Interiano RB, Zhou J, et al. Targeting Histone Demethylases in MYC-Driven Neuroblastomas with Ciclopirox. Cancer research 2017;77:4626–38.

19. Jiang J, Ganesh T, Du Y, Thepchatri P, Rojas A, Lewis I, et al. Neuroprotection by selective allosteric potentiators of the EP2 prostaglandin receptor. Proceedings of the National Academy of Sciences of the United States of America 2010;107:2307–12.

20. Jiang J, Van TM, Ganesh T, Dingledine R. Discovery of 2-Piperidinyl Phenyl Benzamides and Trisubstituted Pyrimidines as Positive Allosteric Modulators of the Prostaglandin Receptor EP2. ACS chemical neuroscience 2018;9:699–707.

21. Sung YM, He G, Fischer SM. Lack of expression of the EP2 but not EP3 receptor for prostaglandin E2 results in suppression of skin tumor development. Cancer research 2005;65:9304–11.

22. Ma X, Aoki T, Tsuruyama T, Narumiya S. Definition of Prostaglandin E2-EP2 Signals in the Colon Tumor Microenvironment That Amplify Inflammation and Tumor Growth. Cancer research 2015;75:2822–32.

23. Merz C, von Massenhausen A, Queisser A, Vogel W, Andren O, Kirfel J, et al. IL-6 Overexpression in ERG-Positive Prostate Cancer Is Mediated by Prostaglandin Receptor EP2. The American journal of pathology 2016;186:974–84.

24. Aoki T, Narumiya S. Prostaglandin E2-EP2 signaling as a node of chronic inflammation in the colon tumor microenvironment. Inflammation and regeneration 2017;37:4.

25. Trigg RM, Turner SD. ALK in Neuroblastoma: Biological and Therapeutic Implications. Cancers 2018;10

26. Middlemas DS, Kihl BK, Zhou J, Zhu X. Brain-derived neurotrophic factor promotes survival and chemoprotection of human neuroblastoma cells. J Biol Chem 1999;274:16451–60.

27. Metelitsa LS, Wu HW, Wang H, Yang Y, Warsi Z, Asgharzadeh S, et al. Natural killer T cells infiltrate neuroblastomas expressing the chemokine CCL2. The Journal of experimental medicine 2004;199:1213–21.

28. Webb MW, Sun J, Sheard MA, Liu WY, Wu HW, Jackson JR, et al. Colony stimulating factor 1 receptor blockade improves the efficacy of chemotherapy against human neuroblastoma in the absence of T lymphocytes. International journal of cancer 2018;143:1483–93.

29. Nevo I, Sagi-Assif O, Meshel T, Ben-Baruch A, Johrer K, Greil R, et al. The involvement of the fractalkine receptor in the transmigration of neuroblastoma cells through bone-marrow endothelial cells. Cancer letters 2009;273:127–39.

30. Hashimoto O, Yoshida M, Koma Y, Yanai T, Hasegawa D, Kosaka Y, et al. Collaboration of cancer-associated fibroblasts and tumour-associated macrophages for neuroblastoma development. The Journal of pathology 2016;240:211–23.

31. Liberman J, Sartelet H, Flahaut M, Muhlethaler-Mottet A, Coulon A, Nyalendo C, et al. Involvement of the CXCR7/CXCR4/CXCL12 axis in the malignant progression of human neuroblastoma. PloS one 2012;7:e43665.

32. Ho R, Minturn JE, Hishiki T, Zhao H, Wang Q, Cnaan A, et al. Proliferation of human neuroblastomas mediated by the epidermal growth factor receptor. Cancer research 2005;65:9868–75.

33. Elaraj DM, Weinreich DM, Varghese S, Puhlmann M, Hewitt SM, Carroll NM, et al. The role of interleukin 1 in growth and metastasis of human cancer xenografts. Clinical cancer research: an official journal of the American Association for Cancer Research 2006;12:1088–96.

34. Pistoia V, Bianchi G, Borgonovo G, Raffaghello L. Cytokines in neuroblastoma: from pathogenesis to treatment. Immunotherapy 2011;3:895–907.

35. Ara T, Nakata R, Sheard MA, Shimada H, Buettner R, Groshen SG, et al. Critical role of STAT3 in IL-6-mediated drug resistance in human neuroblastoma. Cancer research 2013;73:3852–64.

36. Craig BT, Rellinger EJ, Guo Y, Qiao J, Chung DH. Growth Differentiation Factor 15 is a Novel Regulator of the Stem Cell-Like Phenotype in Neuroblastoma. Journal of the American College of Surgeons 2016;223:S87.

37. Sugiura Y, Shimada H, Seeger RC, Laug WE, DeClerck YA. Matrix metalloproteinases-2 and −9 are expressed in human neuroblastoma: contribution of stromal cells to their production and correlation with metastasis. Cancer research 1998;58:2209–16.

38. Nyalendo C, Sartelet H, Barrette S, Ohta S, Gingras D, Beliveau R. Identification of membrane-type 1 matrix metalloproteinase tyrosine phosphorylation in association with neuroblastoma progression. BMC cancer 2009;9:422.

39. Pezzolo A, Parodi F, Corrias MV, Cinti R, Gambini C, Pistoia V. Tumor origin of endothelial cells in human neuroblastoma. Journal of clinical oncology: official journal of the American Society of Clinical Oncology 2007;25:376–83.

40. Sasaki H, Sato Y, Kondo S, Fukai I, Kiriyama M, Yamakawa Y, et al. Expression of the periostin mRNA level in neuroblastoma. Journal of pediatric surgery 2002;37:1293–7.

41. Morra L, Moch H. Periostin expression and epithelial-mesenchymal transition in cancer: a review and an update. Virchows Archiv: an international journal of pathology 2011;459:465–75.

42. Tummler C, Snapkov I, Wickstrom M, Moens U, Ljungblad L, Maria Elfman LH, et al. Inhibition of chemerin/CMKLR1 axis in neuroblastoma cells reduces clonogenicity and cell viability in vitro and impairs tumor growth in vivo. Oncotarget 2017;8:95135–51.

43. Hadjidaniel MD, Muthugounder S, Hung LT, Sheard MA, Shirinbak S, Chan RY, et al. Tumor-associated macrophages promote neuroblastoma via STAT3 phosphorylation and up-regulation of c-MYC. Oncotarget 2017;8:91516–29.

44. Castriconi R, Dondero A, Bellora F, Moretta L, Castellano A, Locatelli F, et al. Neuroblastoma-derived TGF-beta1 modulates the chemokine receptor repertoire of human resting NK cells. Journal of immunology (Baltimore, Md: 1950) 2013;190:5321–8.

45. Tran HC, Wan Z, Sheard MA, Sun J, Jackson JR, Malvar J, et al. TGFbetaR1 Blockade with Galunisertib (LY2157299) Enhances Anti-Neuroblastoma Activity of the Anti-GD2 Antibody Dinutuximab (ch14.18) with Natural Killer Cells. Clinical cancer research: an official journal of the American Association for Cancer Research 2017;23:804–13.

46. Jakovljevic G, Culic S, Stepan J, Bonevski A, Seiwerth S. Vascular endothelial growth factor in children with neuroblastoma: a retrospective analysis. Journal of experimental & clinical cancer research: CR 2009;28:143.

47. Salcedo R, Hixon JA, Stauffer JK, Jalah R, Brooks AD, Khan T, et al. Immunologic and therapeutic synergy of IL-27 and IL-2: enhancement of T cell sensitization, tumor-specific CTL reactivity and complete regression of disseminated neuroblastoma metastases in the liver and bone marrow. Journal of immunology (Baltimore, Md: 1950) 2009;182:4328–38.

48. Alvarez S, Blanco A, Fresno M, Munoz-Fernandez MA. TNF-alpha contributes to caspase-3 independent apoptosis in neuroblastoma cells: role of NFAT. PloS one 2011;6:e16100.

49. Zimmerman MW, Liu Y, He S, Durbin AD, Abraham BJ, Easton J, et al. MYC Drives a Subset of High-Risk Pediatric Neuroblastomas and Is Activated through Mechanisms Including Enhancer Hijacking and Focal Enhancer Amplification. Cancer discovery 2018;8:320–35.

50. O’Brien R, Tran SL, Maritz MF, Liu B, Kong CF, Purgato S, et al. MYC-Driven Neuroblastomas Are Addicted to a Telomerase-Independent Function of Dyskerin. Cancer research 2016;76:3604–17.

51. af Forselles KJ, Root J, Clarke T, Davey D, Aughton K, Dack K, et al. In vitro and in vivo characterization of PF-04418948, a novel, potent and selective prostaglandin EP(2) receptor antagonist. British journal of pharmacology 2011;164:1847–56.

52. Jiang J, Ganesh T, Du Y, Quan Y, Serrano G, Qui M, et al. Small molecule antagonist reveals seizure-induced mediation of neuronal injury by prostaglandin E2 receptor subtype EP2. Proceedings of the National Academy of Sciences of the United States of America 2012;109:3149–54.

53. Ganesh T, Jiang J, Shashidharamurthy R, Dingledine R. Discovery and characterization of carbamothioylacrylamides as EP2 selective antagonists. ACS Med Chem Lett 2013;4:616–21.

54. Jiang J, Quan Y, Ganesh T, Pouliot WA, Dudek FE, Dingledine R. Inhibition of the prostaglandin receptor EP2 following status epilepticus reduces delayed mortality and brain inflammation. Proceedings of the National Academy of Sciences of the United States of America 2013;110:3591–6.

55. Dey A, Kang X, Qiu J, Du Y, Jiang J. Anti-Inflammatory Small Molecules To Treat Seizures and Epilepsy: From Bench to Bedside. Trends Pharmacol Sci 2016;37:463–84.

56. Kang X, Qiu J, Li Q, Bell KA, Du Y, Jung DW, et al. Cyclooxygenase-2 contributes to oxidopamine-mediated neuronal inflammation and injury via the prostaglandin E2 receptor EP2 subtype. Scientific reports 2017;7:9459.

57. Yu Y, Nguyen DT, Jiang J. G protein-coupled receptors in acquired epilepsy: Druggability and translatability. Progress in neurobiology 2019;183:101682.

58. Jiang J, Yu Y, Kinjo ER, Du Y, Nguyen HP, Dingledine R. Suppressing pro-inflammatory prostaglandin signaling attenuates excitotoxicity-associated neuronal inflammation and injury. Neuropharmacology 2019;149:149–60.

59. Du Y, Kemper T, Qiu J, Jiang J. Defining the therapeutic time window for suppressing the inflammatory prostaglandin E2 signaling after status epilepticus. Expert Rev Neurother 2016;16:123–30.

60. Savonenko A, Munoz P, Melnikova T, Wang Q, Liang X, Breyer RM, et al. Impaired cognition, sensorimotor gating, and hippocampal long-term depression in mice lacking the prostaglandin E2 EP2 receptor. Exp Neurol 2009;217:63–73.

61. Yang H, Zhang J, Breyer RM, Chen C. Altered hippocampal long-term synaptic plasticity in mice deficient in the PGE2 EP2 receptor. J Neurochem 2009;108:295–304.

62. Rossler J, Taylor M, Geoerger B, Farace F, Lagodny J, Peschka-Suss R, et al. Angiogenesis as a target in neuroblastoma. European journal of cancer (Oxford, England: 1990) 2008;44:1645–56.

63. DeLisser HM, Christofidou-Solomidou M, Strieter RM, Burdick MD, Robinson CS, Wexler RS, et al. Involvement of endothelial PECAM-1/CD31 in angiogenesis. The American journal of pathology 1997;151:671–7.

64. Grosser T, Yu Y, Fitzgerald GA. Emotion recollected in tranquility: lessons learned from the COX-2 saga. Annual review of medicine 2010;61:17–33.

65. O’Callaghan G, Houston A. Prostaglandin E2 and the EP receptors in malignancy: possible therapeutic targets? British journal of pharmacology 2015;172:5239–50.

66. Yu Y, Ricciotti E, Scalia R, Tang SY, Grant G, Yu Z, et al. Vascular COX-2 modulates blood pressure and thrombosis in mice. Science translational medicine 2012;4:132ra54.

67. Jiang J, Dingledine R. Prostaglandin receptor EP2 in the crosshairs of anti-inflammation, anti-cancer, and neuroprotection. Trends Pharmacol Sci 2013;34:413–23.

68. Okuyama T, Ishihara S, Sato H, Rumi MA, Kawashima K, Miyaoka Y, et al. Activation of prostaglandin E2-receptor EP2 and EP4 pathways induces growth inhibition in human gastric carcinoma cell lines. The Journal of laboratory and clinical medicine 2002;140:92–102.

69. Spinella F, Rosano L, Di Castro V, Natali PG, Bagnato A. Endothelin-1-induced prostaglandin E2-EP2, EP4 signaling regulates vascular endothelial growth factor production and ovarian carcinoma cell invasion. J Biol Chem 2004;279:46700–5.

70. Jain S, Chakraborty G, Raja R, Kale S, Kundu GC. Prostaglandin E2 regulates tumor angiogenesis in prostate cancer. Cancer research 2008;68:7750–9.

71. Rundhaug JE, Simper MS, Surh I, Fischer SM. The role of the EP receptors for prostaglandin E2 in skin and skin cancer. Cancer metastasis reviews 2011;30:465–80.

72. Ma X, Kundu N, Collin PD, Goloubeva O, Fulton AM. Frondoside A inhibits breast cancer metastasis and antagonizes prostaglandin E receptors EP4 and EP2. Breast cancer research and treatment 2012;132:1001–8.

73. Hsu HH, Lin YM, Shen CY, Shibu MA, Li SY, Chang SH, et al. Prostaglandin E2-Induced COX-2 Expressions via EP2 and EP4 Signaling Pathways in Human LoVo Colon Cancer Cells. International journal of molecular sciences 2017;18

74. Wang J, Zhang L, Kang D, Yang D, Tang Y. Activation of PGE2/EP2 and PGE2/EP4 signaling pathways positively regulate the level of PD-1 in infiltrating CD8(+) T cells in patients with lung cancer. Oncology letters 2018;15:552–8.

75. Donnini S, Finetti F, Solito R, Terzuoli E, Sacchetti A, Morbidelli L, et al. EP2 prostanoid receptor promotes squamous cell carcinoma growth through epidermal growth factor receptor transactivation and iNOS and ERK1/2 pathways. FASEB journal: official publication of the Federation of American Societies for Experimental Biology 2007;21:2418–30.

76. Kamiyama M, Pozzi A, Yang L, DeBusk LM, Breyer RM, Lin PC. EP2, a receptor for PGE2, regulates tumor angiogenesis through direct effects on endothelial cell motility and survival. Oncogene 2006;25:7019–28.

77. Finetti F, Solito R, Morbidelli L, Giachetti A, Ziche M, Donnini S. Prostaglandin E2 regulates angiogenesis via activation of fibroblast growth factor receptor-1. J Biol Chem 2008;283:2139–46.

78. Trau HA, Brannstrom M, Curry TE, Jr., Duffy DM. Prostaglandin E2 and vascular endothelial growth factor A mediate angiogenesis of human ovarian follicular endothelial cells. Human reproduction (Oxford, England) 2016;31:436–44.

79. O’Brien AJ, Fullerton JN, Massey KA, Auld G, Sewell G, James S, et al. Immunosuppression in acutely decompensated cirrhosis is mediated by prostaglandin E2. Nature medicine 2014;20:518–23.

80. Quan Y, Jiang J, Dingledine R. EP2 receptor signaling pathways regulate classical activation of microglia. J Biol Chem 2013;288:9293–302.

